# Aquaporin-9 and aquaporin-10 but not aquaporin-3 confer susceptibility to dimethylarsinic acid genotoxicity in human cells

**DOI:** 10.64898/2026.09.20.752519

**Authors:** Shiwei Yin, Bingru Feng, Maryam Vaziripour, Jun Xia

## Abstract

Human metabolism converts inorganic arsenic to the pentavalent methylated species MMA(V) and DMA(V), the forms most people excrete, and the forms long read as the end of a detoxification pathway. Whether a transporter sets how much of these metabolites reaches the genome has not been tested in a mammalian cell. We expressed human AQP3, AQP7, AQP9 or AQP10 in HEK293T and MRC5-SV40 cells and measured γH2AX by flow cytometry across dose series of As(V), MMA(V) and DMA(V), pairing every aquaporin with a GFP-Tubulin control and an untransfected mock acquired in the same run. As(V) was inactive in HEK293T cells and only weakly active in MRC5-SV40 cells to 20 µM, and both methylated species damaged DNA only in the millimolar range, DMA(V) being the more potent of the two in both cell lines. Against that weak baseline, AQP9 and AQP10 raised DMA(V)-induced γH2AX in HEK293T cells by roughly 17 percentage points over the matched control, more than doubling the damage the same exposure produced in control cells, whereas AQP3 and AQP7 changed it not at all. AQP9 alone remained active with MMA(V). The ranking held in MRC5-SV40 fibroblasts at one-sixth the size, and within single wells the damage rose with the amount of AQP9 a cell carried while the control was flat. Aquaglyceroporins therefore discriminate among arsenic species, and AQP9 and AQP10 turn a weakly genotoxic metabolite into a substantially more genotoxic one.

**Highlights:** • AQP9 and AQP10 more than double DMA(V)-induced DNA damage in human cells.

• AQP3 and AQP7 show no such effect: permeability differs between paralogues.

• AQP9 alone remains active with MMA(V); AQP10 does not.

• Damage rises with the amount of AQP9 a single cell carries.

• Genotoxic potency depends on the transporter complement of the exposed cell.

**Impact statement:** Arsenic risk assessment treats the pentavalent methylated metabolites as detoxification products, and on their own they are weak genotoxins. We show that this weakness is conditional on the transporter complement of the exposed cell. AQP9 and AQP10 more than double the DNA damage a given DMA(V) exposure produces in human cells, while AQP3, which confers arsenite susceptibility in the same laboratory, does not. An aquaglyceroporin and a poorly toxic arsenical therefore act together to produce damage that neither produces alone, so the genotoxic potency of an arsenical cannot be read from the species and the dose without knowing which channels admit it. Uptake capacity, which differs between tissues and between people, belongs alongside metabolism among the factors that set internal dose and susceptibility.

## 1. Introduction

Inorganic arsenic (iAs) is among the most widespread environmental carcinogens and is classified by IARC as carcinogenic to humans^1^. Chronic exposure through groundwater affects more than 200 million people and causes cancer of the lung, urinary bladder and skin^2,3^. Most of an ingested dose does not remain inorganic: human metabolism oxidatively methylates iAs, and the pentavalent species MMA(V) and DMA(V) are the dominant urinary forms^4^.

Methylation was long read as detoxification, and the pentavalent metabolites are indeed far less potent than their trivalent counterparts^5,6^. That reading is incomplete. DMA(V) is a complete urinary bladder carcinogen in the rat^7,8^, and whether the pentavalent species damage DNA in human cells remains contested: an alkaline comet screen in TK6 lymphoblastoid cells found MMA(V) and DMA(V) inactive^9^, whereas millimolar DMA(V) induced strand breaks in the human alveolar line L-132 through a peroxyl-radical mechanism^10^. The positive evidence otherwise comes from rodents in vivo, where DMA(V) produces oxidative DNA damage and strand breaks^10,11^ but is at most marginally mutagenic^12^.

Uptake resolves much of that disagreement, because internal rather than medium concentration sets genotoxic potency and arsenic species differ by orders of magnitude in how efficiently they cross a membrane^13,14^. At physiological pH trivalent arsenic exists largely as uncharged As(OH)_3_ and enters through the aquaglyceroporins^15–17^, a subfamily whose members differ in the solutes they carry^18^. The methylated species use the same route: aquaglyceroporins conduct MMA(III)^19^, human AQP9 in Xenopus oocytes transports both MMA(V) and DMA(V)^20^, and a bacterial aquaglyceroporin conducts trivalent and pentavalent methylarsenicals alike^21^.

Those studies measure flux across a membrane. Whether that flux is large enough, in a human cell with its own efflux and metabolism, to change how much DNA damage an exposure produces has not been determined.

Two members of the subfamily are poorly served by the existing literature. AQP10 has been tested only with inorganic arsenic and never with a methylated arsenical, and no genotoxic endpoint has been reported for it in any system: human AQP10 conducted little or no As(OH)_3_ in oocytes^22^, silencing it in Caco-2 cells altered As(III) uptake^23^, and the remaining comparable work is in zebrafish^24^. Mouse Aqp10 is a pseudogene^25^, so the channel is absent from the rodent models on which arsenic toxicology rests, although human AQP10 is a functional glycerol channel^26^. AQP3 is disputed: it confers arsenite susceptibility in human lung fibroblasts ^27^ and reduced AQP3 expression accompanies lower arsenic accumulation in an arsenite-resistant lung adenocarcinoma subline^28^, yet a killifish ortholog conducts water, glycerol and urea but not arsenite^29^. Here we expressed all four human aquaglyceroporins in two human cell lines and show that AQP9 and AQP10 confer susceptibility to DMA(V)-induced DNA damage while AQP3 and AQP7 do not, that AQP9 alone remains active with MMA(V), and that the damage a cell takes rises with the amount of channel it carries. Aquaglyceroporin expression therefore lowers the concentration at which the dominant human arsenic metabolite becomes genotoxic.

## 2. Materials and methods

### 2.1 AQP overexpression constructs and cloning

Gateway entry clones for AQP3, AQP7, AQP9 and AQP10 were synthesized and sequence-verified in pDONR vectors and recombined into the N-terminal GFP destination vector pcDNA6.2/N-EmGFP-DEST with Gateway LR Clonase II (Thermo Fisher, 11791020) following the manufacturer’s protocol. Each LR reaction combined 50 to 150 ng of entry clone with 150 ng of destination vector in TE buffer, pH 8.0, to a final volume of 8 µL, received 2 µL of LR Clonase II, was incubated at 25 °C for 1 h and was terminated with 1 µL of proteinase K (Thermo Fisher, EO0491) at 37 °C for 10 min. Reactions were transformed into One Shot OmniMAX 2-T1R competent cells (Thermo Fisher, C854003) and plated on LB agar containing 100 µg/mL ampicillin (Sigma-Aldrich, A9518); single colonies were expanded and plasmid DNA recovered with a QIAprep spin miniprep kit (Qiagen, 27106). Every construct was verified by BsrGI-HF digestion (New England Biolabs, R3575S) and by whole plasmid sequencing. GFP-Tubulin was the transfection control throughout and untransfected mock the gating control. GFP-Tubulin controls for transfection, for GFP expression and for the fluorescent tag.

### 2.2 cell culture

HEK293T and MRC5-SV40 (SV40-immortalised human lung fibroblasts) are the stocks used previously in this laboratory^30^. Both were cultured in Dulbecco’s modified Eagle’s medium (DMEM; Invitrogen, 11965-092) supplemented with 10% fetal bovine serum, 1x 2 mM L-glutamine, penicillin and streptomycin (Gibco), maintained at 37 °C in 5% CO_2_ and passaged before confluence. Lines were authenticated by short tandem repeat profiling (ATCC) and tested routinely for mycoplasma. Both lines carry SV40 large T antigen and therefore have compromised checkpoint signaling.

### 2.3 Arsenical exposure and transfection

Cells were seeded at 5 × 10^5^ per well in 6-well plates and the medium was replaced with fresh serum-containing DMEM 30 min before transfection. For each well, 1 µg of plasmid DNA was diluted in 40 µL of serum-free DMEM and 2.5 µL of GenJet In Vitro DNA Transfection Reagent Ver. II (SignaGen, SL100488) was diluted separately in 40 µL of serum-free DMEM; each was held for 5 min, the two were combined and held for a further 10 min at room temperature, and the resulting 80 µL complex was added dropwise to the well.

Arsenicals were added to the medium immediately after transfection and again at the same nominal concentration when the medium was replaced the following day, so the exposure ran for 72 h with one restoration, and cells were harvested 72 h after transfection. The three species were sodium arsenate dibasic heptahydrate for As(V) (Sigma, A6756-50G), disodium methyl arsonate hexahydrate for MMA(V) (Santa Cruz Biotechnology, sc-257380) and cacodylic acid for DMA(V) (Sigma, 20835-50G-F). Doses are given throughout as the molar concentration of the arsenical in the medium, and vehicle denotes the matched 0 µM or 0 mM well of the same construct from the same replicate, which received medium without arsenical. Dose-finding series covered As(V) at 0, 2, 5, 10 and 20 µM, DMA(V) at 0, 0.5, 1 and 2 mM and MMA(V) at 0, 5, 10 and 20 mM in untransfected cells. The transfected panel used DMA(V) at 0, 1 and 2 mM and MMA(V) at 0, 0.5, 1, 2.5 and 5 mM in HEK293T cells, and DMA(V) at 0, 0.25 and 0.5 mM and MMA(V) at 0, 0.5 and 1 mM in MRC5-SV40 cells.

### 2.4 Flow cytometry and gating

Phosphorylation of histone H2AX on Ser139 (γH2AX) spreads over megabase chromatin domains flanking a DNA double-strand break, so the proportion of cells with high γH2AX signal reports the fraction of the population mounting a double-strand-break response^31,32^. γH2AX was measured by indirect immunofluorescence on fixed, permeabilised intact cells, as described previously for these lines^30^. Cells were harvested by trypsinisation and approximately 1 × 10^6^ cells per tube were carried into staining, pelleted at 1,000 × g for 5 min at 4 °C, fixed in freshly prepared 2% (w/v) formaldehyde in PBS for 15 min on ice, permeabilised in 0.05% Triton X-100 for 15 min on ice and blocked in 5% BSA/PBS (Cell Signaling Technology, 9998S) for 30 min on ice. Cells were stained with anti-phospho-histone H2A.X (Ser139) clone JBW301 (1:750; Sigma-Aldrich, 05-636-25UG) for 1 h at room temperature, washed, and incubated with goat anti-mouse IgG (H+L) Alexa Fluor 647 (1:1,000; Invitrogen, A-21235) for 1 h at room temperature in the dark. Every experiment carried at least an untransfected, antibody-stained mock well, which supplied that replicate’s gating thresholds. Data were acquired on a Bio-Rad YETI/ZE5 cytometer. Gating was applied identically to all 288 files and is shown in Supplementary Fig. 1A–D: a robust 95% Mahalanobis contour on log FSC-H × log SSC-H defined cells, a ±3 MAD window on log FSC-H − log FSC-A defined singlets, and GFP-positive cells were those above the 99.9th percentile of the same replicate’s untransfected mock. Height rather than area channels were used because FSC-A saturated in up to 9% and SSC-A in up to 35% of events, whereas the height channels did not.

The γH2AX-high gate was set at the top 0.5%, 1%, 2% and 5% of each replicate’s own untreated mock (Supplementary Fig. 1E, F). Every contrast was computed at all four settings and the top-1% gate is reported in the main text; the gate sweep is shown in every figure so that no result depends on one threshold. Untransfected mock carries about 0.1% GFP-positive events by design, so mock is read on its GFP-negative population throughout.

### 2.5 Endpoint, expression-DNA damage analysis and statistics

The endpoint is vehicle-subtracted induction: the γH2AX-high percentage of a well minus that of the same construct’s 0 mM well in the same replicate. Aquaporins were compared with the GFP-Tubulin well beside them in the same replicate at the same dose, and GFP-Tubulin with untransfected mock, by one-tailed paired t-test on the replicate × dose pairs, pooled over treated doses. Per-dose values are given in Supplementary Fig. 3A-D. Analyses were performed in Python 3 with NumPy, pandas and SciPy; code and the per-well table are available as described under Data availability.

For the expression–response analysis, GFP-positive cells of each well were divided into fifths by GFP intensity and the γH2AX-high percentage computed in each fifth. Each construct’s treated value was corrected fifth by fifth against its own vehicle well in the same replicate. The slope is the ordinary least-squares regression of that corrected value on fifth number (1–5), in percentage points per fifth, given as mean ± SEM across replicates; the dimmest-to-brightest difference is the corrected value in fifth 5 minus that in fifth 1.

## 3. Results

### 3.1 As(V) is inactive or weakly active to 20 µM; DMA(V) and MMA(V) act in the millimolar range

To place the three species on one scale before any transporter was introduced, we exposed untransfected cells of both lines to each arsenical across a dose-finding range (Fig. 1A, B). As(V) did not raise the γH2AX-high fraction in HEK293T cells to 20 µM (0.015 percentage points per µM, R^2^ = 0.20, P = 0.14) and produced a shallow response in MRC5-SV40 cells that reached 1.95% at 20 µM from 0.36% in vehicle (0.086 points per µM, R^2^ = 0.90, P < 0.0001).

**Figure 1.**
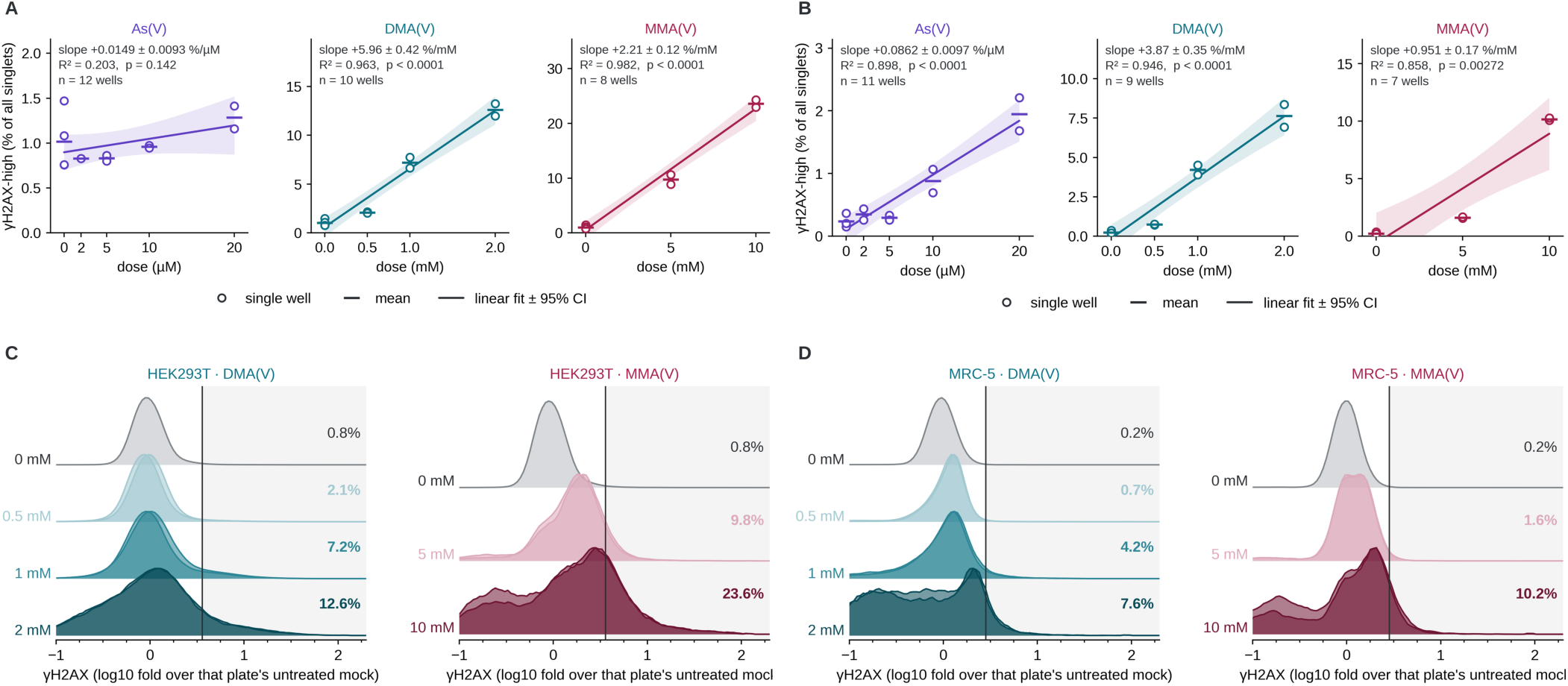
Arsenic speciation sets the active concentration range in untransfected cells. (**A**) HEK293T and (**B**) MRC5-SV40 mock dose-finding series for As(V), DMA(V) and MMA(V). Open circles, single wells; bars, per-dose means; line, ordinary least-squares regression of γH2AX-high on dose with its 95% confidence band; slope ± SE, R^2^ and the two-sided P for slope = 0 given in each panel. Mock is untransfected, so the readout is the whole singlet population. MMA(V) 20 mM is excluded: γH2AX falls there while event recovery collapses. (**C**, **D**) γH2AX distributions across the same ladders in HEK293T and MRC5-SV40 cells, one ridge per dose, normalised to the mode; vertical line and shading, the top-1% gate of that replicate’s untreated mock; percentages, γH2AX-high at each dose. The sub-vehicle shoulder at the top dose of each ladder is the dying fraction.

Both methylated species were active only in the millimolar range. DMA(V) rose from 0.76% in vehicle to 2.09%, 7.20% and 12.60% across 0.5, 1 and 2 mM in HEK293T cells, and from 0.15% to 0.75%, 4.21% and 7.64% in MRC5-SV40 cells. MMA(V) required higher concentrations, reaching 9.77% at 5 mM and 23.62% at 10 mM in HEK293T cells and 1.62% and 10.17% at the same doses in MRC5-SV40 cells.

DMA(V) was 2.7-fold more potent than MMA(V) per mole in HEK293T cells and 3.7-fold in MRC5-SV40 cells, reaching five percentage points above vehicle at 0.86 against 2.78 mM and at 1.27 against 7.09 mM. The whole γH2AX distribution shifted with dose rather than a tail crossing a threshold (Fig. 1C, D; the same distributions as conventional overlays in Supplementary Fig. 2).

MMA(V) inverted above its usable window. At 20 mM the γH2AX-high fraction fell to 4.63% in HEK293T and 5.62% in MRC5-SV40 cells, a sub-vehicle population appeared below the main peak, and those wells carry the fewest events in the dataset, 3,894 to 6,774 against 0.2 to 0.9 million in vehicle. A damage marker that falls while event recovery falls with it reports cell death, so 20 mM is excluded from every fit and contrast and the MMA(V) response is not described as biphasic.

### 3.2 AQP9 and AQP10 confer susceptibility to DMA(V)-induced DNA damage

To determine whether aquaglyceroporin expression changes the damage a given DMA(V) exposure produces, we compared each aquaporin with the GFP-Tubulin well beside it in the same replicate at the same dose (Fig. 2A). AQP9 raised DMA(V)-induced γH2AX in HEK293T cells by 16.96 ± 2.92 percentage points (P = 0.005, positive in four of four pairs) and AQP10 by 16.54 ± 4.62 points (P = 0.019, four of four). AQP3 gave −0.11 ± 1.60 points (P = 0.52) and AQP7 −4.22 ± 2.26 points (P = 0.92). Both active differences are large against the underlying response, which reaches 12.60% in untransfected cells at the top dose, and both widen rather than narrow as the γH2AX gate is relaxed from the top 0.5% to the top 5% of control, the behaviour of a shifted population rather than of a few cells crossing a threshold (Fig. 2A, columns). Representative single-well distributions at 2 mM are shown in Fig. 2B: the AQP9 and AQP10 curves clear the gate as whole populations, while the AQP3 and AQP7 curves sit on the GFP-Tubulin curve.

**Figure 2.**
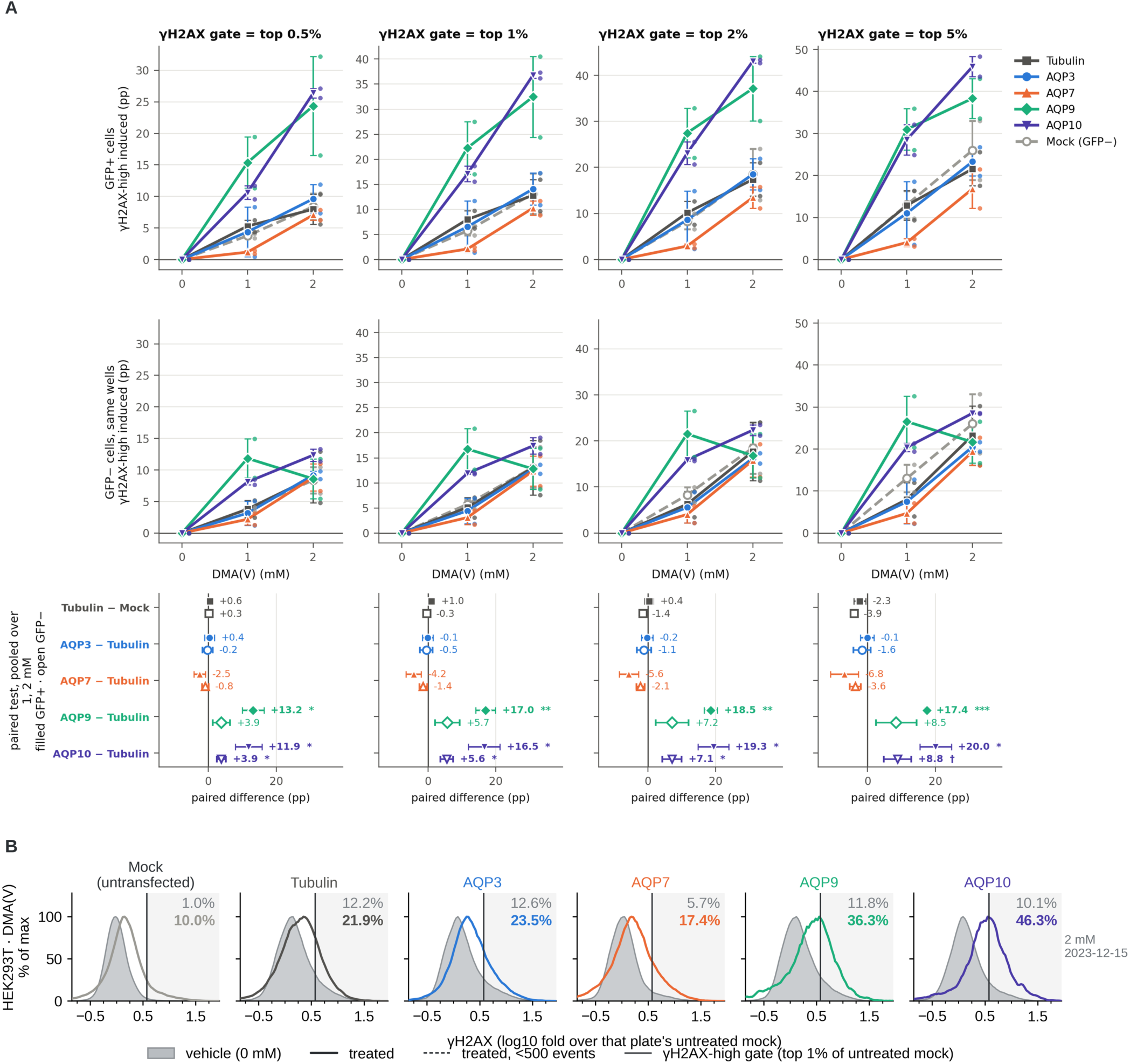
AQP9 and AQP10 confer susceptibility to DMA(V) in HEK293T cells. (**A**) Vehicle-subtracted γH2AX induction across the DMA(V) dose range, two replicates. Columns are the four γH2AX gates. Row 1, GFP-positive cells; row 2, the GFP-negative cells of the same wells; row 3, paired differences pooled over treated doses, filled symbols GFP-positive and open symbols GFP-negative, mean ± SEM, one-tailed paired t-test on replicate × dose pairs. Mock is untransfected and is read on its whole population in both rows. (**B**) Representative single-well γH2AX distributions for every construct at 2 mM DMA(V): vehicle filled, treated in colour, vertical line the top-1% gate, percentages γH2AX-high with vehicle above treated; the replicate is identified in the panel. † P < 0.10, * P < 0.05, ** P < 0.01, *** P < 0.001.

Transfection alone did not account for this. GFP-Tubulin cells differed from untransfected mock by 0.98 ± 0.90 points for DMA(V) (P = 0.18), so the aquaporin differences sit on an unchanged baseline.

### 3.3 With MMA(V), AQP9 alone remains active and transfection itself sensitises

Repeating the comparison for MMA(V) in HEK293T cells over eight replicate × dose pairs separated the two metabolites (Fig. 3A, with representative distributions in Fig. 3B). AQP9 gave 9.23 ± 4.54 points (P = 0.041), roughly half its DMA(V) value, while AQP10 fell to 2.12 ± 3.82 (P = 0.30), AQP3 to 1.11 ± 2.35 (P = 0.33) and AQP7 to −3.99 ± 1.70 (P = 0.97).

**Figure 3.**
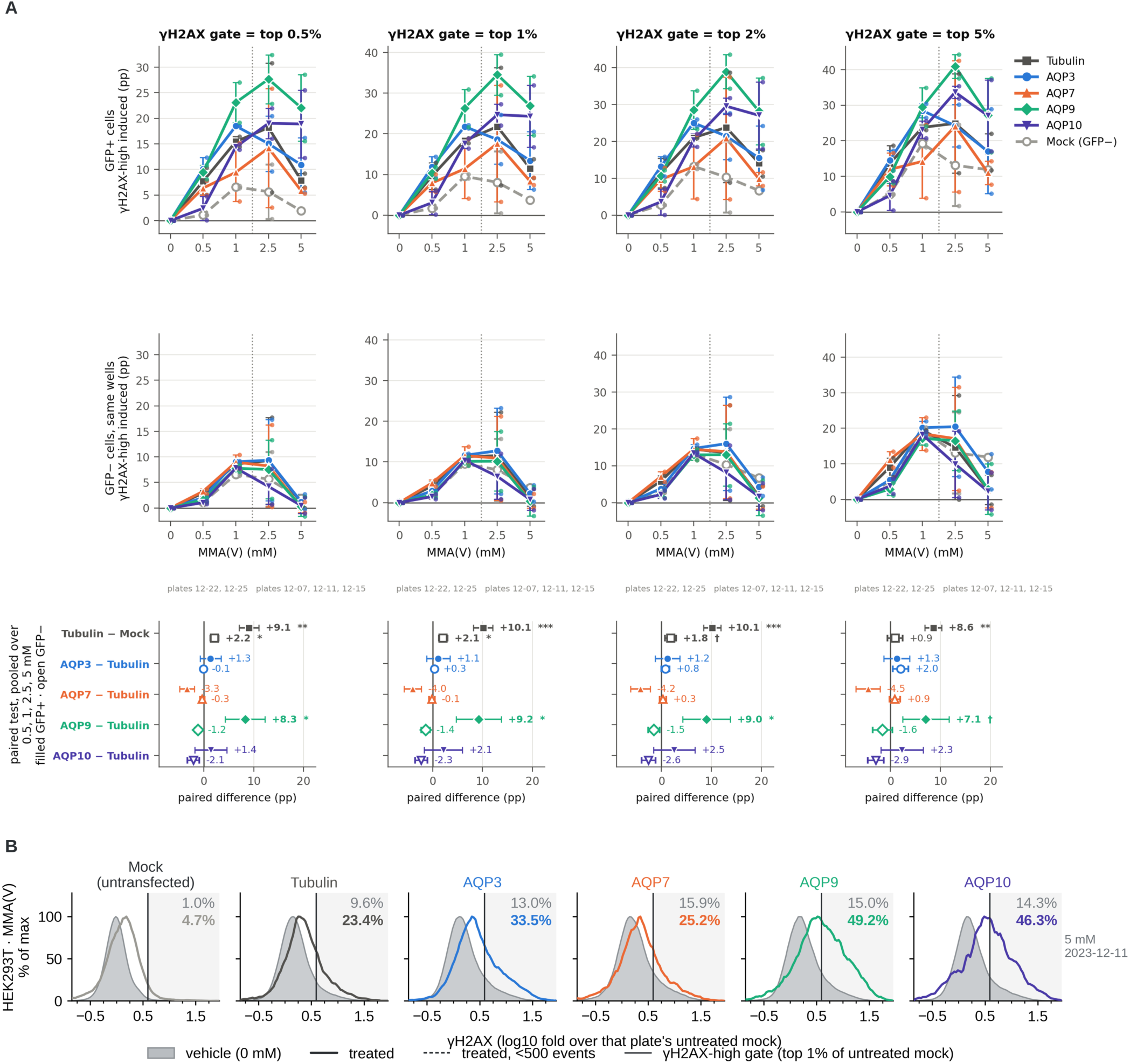
With MMA(V) in HEK293T cells, AQP9 alone remains active and transfection itself sensitizes. (**A**) Vehicle-subtracted γH2AX induction across the MMA(V) dose range, two replicates at 0.5 and 1 mM and three at 2.5 and 5 mM; the dotted vertical rule separates the two replicate sets. (**B**) Representative single-well distributions at 5 mM MMA(V). Layout, gates and statistics as in Fig. 2. The Tubulin-minus-mock contrast in row 3 is the transfection effect described in section 3.3.

Part of that difference lies outside the aquaporins. Carrying a construct at all changed the MMA(V) response: GFP-Tubulin cells took 10.12 ± 1.93 percentage points more damage from MMA(V) than untransfected mock (P = 0.001, seven of seven pairs), against 0.98 ± 0.90 points for DMA(V). MMA(V) uptake therefore responds to membrane state in general, which both raises the comparator every aquaporin is measured against and leaves less headroom above it. The same contrast in MRC5-SV40 cells ran in the same direction at 3.74 ± 2.53 points (P = 0.12). AQP9 therefore increases MMA(V)-induced γH2AX above an already elevated transfected baseline. Because that baseline is the comparator for every construct, the null results for AQP3, AQP7 and AQP10 in this arm constitute weaker evidence of absence than their DMA(V) counterparts.

### 3.4 Untransfected cells in the same wells carry part of the effect

Every well contains transfected and untransfected cells that received the same dose and the same stain, so the GFP-negative population is an internal control for well-level differences. In the HEK293T DMA(V) arm it was not flat. AQP10 raised γH2AX in the GFP-negative cells of its own wells by 5.57 ± 1.87 points over the GFP-negative cells of the Tubulin wells (P = 0.029) and AQP9 by 5.70 ± 3.59 points (P = 0.11), against −0.50 for AQP3 and −1.35 for AQP7 (Fig. 2A, row 2). The same two constructs carried the same signal in MRC5-SV40 cells, AQP9 at 1.45 ± 0.40 points (P = 0.006) and AQP10 at 0.93 ± 0.44 (P = 0.038) (Fig. 4A, row 2), and it was absent from the HEK293T MMA(V) arm, where AQP9 and AQP10 read −1.41 and −2.28 points (Fig. 3A, row 2).

**Figure 4.**
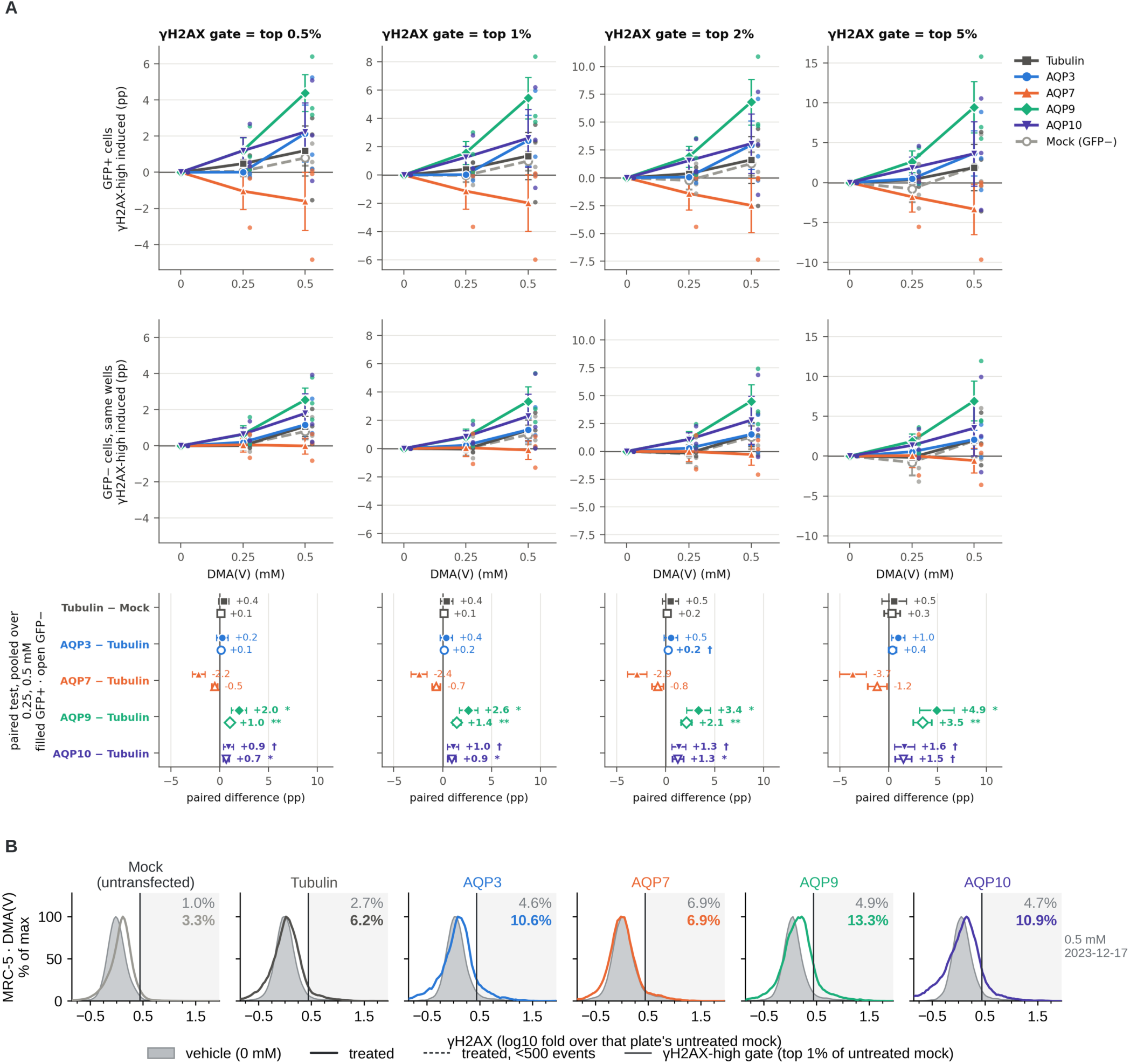
The aquaglyceroporin contribution to DMA(V) damage is smaller in MRC5-SV40 fibroblasts. (**A**) Vehicle-subtracted γH2AX induction across the DMA(V) dose range, three replicates. (**B**) Representative single-well distributions at 0.5 mM DMA(V). Layout, gates and statistics as in Fig. 2.

The effect therefore appears only in the arms and only for the constructs that show a GFP-positive effect, and it is largest at 1 mM DMA(V) (Supplementary Fig. 3A). Three readings are available and this dataset does not separate them: a bystander effect, in which damage initiated in transfected cells is transmitted to their neighbours through a diffusible signal such as reactive oxygen species; spillover of dim GFP-positive cells into the GFP-negative gate; or a well-level difference that the construct produces without the channel acting in the neighbouring cell.

Because the GFP-positive effect is two to three times larger than the GFP-negative one in both DMA(V) arms, the transfected cells carry most of the response whichever reading holds.

### 3.5 The aquaglyceroporin contribution depends on the host cell

In MRC5-SV40 fibroblasts the ranking held and the size fell (Fig. 4A, with representative distributions in Fig. 4B). For DMA(V), AQP9 gave 2.63 ± 0.94 points (P = 0.017, five of six pairs), AQP10 1.05 ± 0.56 (P = 0.058), AQP3 0.36 ± 0.63 (P = 0.30) and AQP7 −2.43 ± 0.82 (P = 0.98). Against 16.96 points for AQP9 in HEK293T cells that is a reduction of roughly six-fold. Both lines carry SV40 large T antigen, so the difference does not follow from p53 status.

The MRC5-SV40 MMA(V) arm supports the same ordering weakly and no contrast in it reached significance (Fig. 5A, B): AQP9 4.82 ± 3.06 points (P = 0.11), AQP3 3.38 ± 2.91, AQP7 1.14 ± 2.28 and AQP10 −2.69 ± 2.12, over four pairs. One well behind those numbers holds 143 GFP-positive events and is the paired comparator for all four aquaporins at 1 mM, so this arm is reported for completeness and no conclusion rests on it.

**Figure 5.**
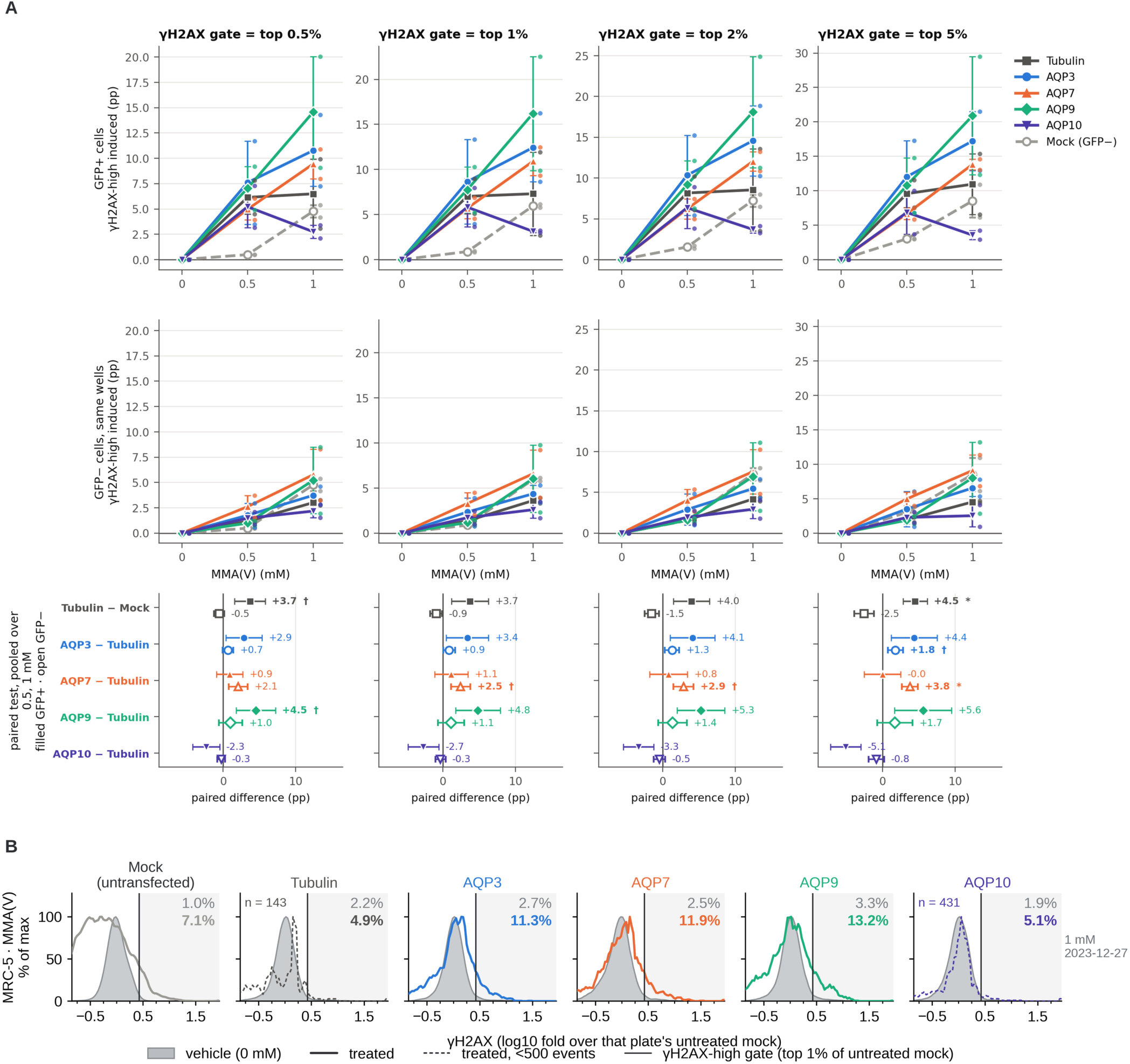
The MRC5-SV40 MMA(V) arm supports the same conclusion. (**A**) Vehicle-subtracted γH2AX induction across the MMA(V) dose range, two treated replicates with a third contributing vehicle wells only. (**B**) Representative single-well distributions at 1 mM MMA(V). Layout, gates and statistics as in Fig. 2. No contrast in A reached significance and one comparator well at 1 mM holds 143 GFP-positive events, so this figure is shown for completeness and no conclusion rests on it.

### 3.6 Damage scales with the amount of aquaporin a cell carries

If the aquaporins act by raising internal dose, cells carrying more channel should take more damage. We split the GFP-positive cells of each well into fifths by GFP intensity and scored γH2AX in each fifth, at the dose in each arm where AQP9 differed most from control (Fig. 6A). γH2AX rose with GFP intensity in every construct, including GFP-Tubulin and including untreated wells, from 4.7% in the dimmest fifth to 23.4% in the brightest of the HEK293T Tubulin vehicle, so brightly transfected cells are more damaged whatever they carry and the comparison must be made against each construct’s own vehicle bin by bin (Fig. 6A).

**Figure 6.**
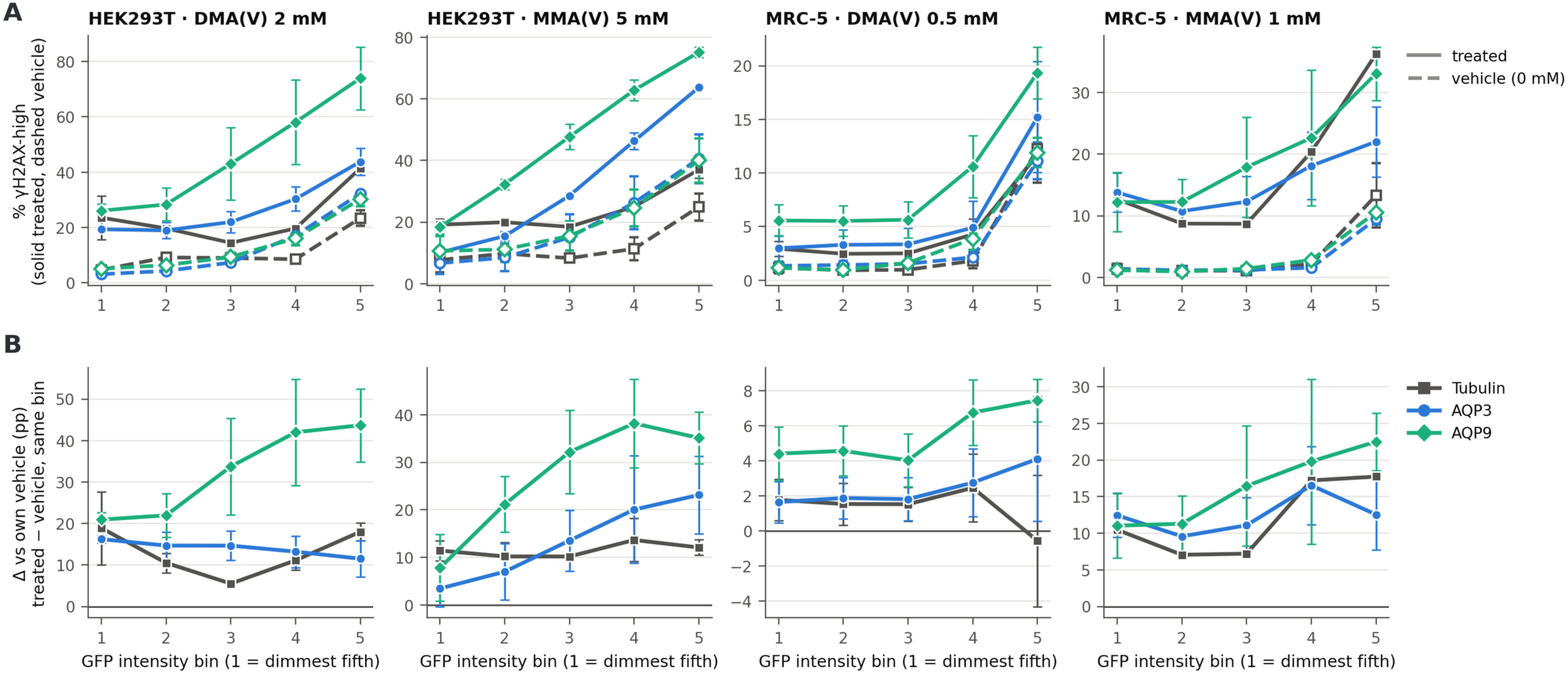
γH2AX-induced damage scales with the amount of aquaporin a cell carries. GFP-positive cells of each well split into fifths by GFP intensity, at the dose in each arm where AQP9 differed most from control; mean ± SEM across replicates, top-1% gate. (**A**) γH2AX-high per fifth for treated (solid) and the same construct’s own vehicle (dashed). (**B**) Each construct minus its own vehicle, fifth by fifth. Columns, the four cell line × arsenical arms. γH2AX rises with GFP intensity in every construct including GFP-Tubulin and in untreated wells, so the vehicle-corrected slope in B is the comparison that isolates the transporter. The GFP intensity spanned by the five fifths is quantified in section 3.7.

After that correction the slope separated the constructs (Fig. 6B). Vehicle-corrected damage rose with AQP9 level in all four arms, at 6.55 ± 2.16, 7.17 ± 0.03, 0.83 ± 0.10 and 3.14 ± 0.64 percentage points per fifth for HEK293T DMA(V), HEK293T MMA(V), MRC5-SV40 DMA(V) and MRC5-SV40 MMA(V), while GFP-Tubulin was flat in three of the four (−0.12, +0.48, −0.37 and +2.47). Between the dimmest and brightest fifths the vehicle-corrected difference for AQP9 was 22.8, 27.3, 3.0 and 11.5 percentage points, against −0.9, +0.7, −2.3 and +7.3 for GFP-Tubulin. The fifths are within-well quantiles rather than matched absolute expression, so this is an expression-response relationship within each well and not a calibrated dose of channel.

### 3.7 Neither transfection efficiency nor expression level accounts for the ranking

Two measurements bear on whether the ranking follows the identity of the channel or the amount of protein each construct made. Transfection efficiency ran against the ranking rather than with it: the median GFP-positive fraction was 35.4% for GFP-Tubulin, 35.3% for AQP9, 33.9% for AQP3, 26.7% for AQP7 and 24.8% for AQP10, so AQP10, the least efficient of the five, produced one of the two largest differences.

GFP intensity gives a semiquantitative read on how much fusion protein the transfected cells carried, and it separates the constructs no better. Taken as the event-weighted mean of the per-fifth median GFP intensities, construct level was 1.41- and 1.81-fold that of GFP-Tubulin for AQP9 in the HEK293T DMA(V) and MMA(V) arms and 1.41- and 1.46-fold in the corresponding MRC5-SV40 arms, against 1.54- and 1.69-fold for AQP3 in HEK293T cells and 0.79- and 0.86- fold in MRC5-SV40 cells. AQP3 therefore reached GFP levels equal to or above AQP9 in the line and the arms where AQP9 was most active, and produced no effect, so the difference between the two is not a difference in how much protein was overproduced.

The same measurement calibrates the GFP fifths of section 3.6 and turns them into an expression ladder. Within a single well the median GFP intensity of the dimmest fifth stood 1.9 to 5.8 times the GFP-positive threshold and that of the brightest fifth 363 to 1,726 times it, a 97- to 543-fold range across the four arms with a median of 415-fold, so the five gates span more than two orders of magnitude of construct level in cells that shared a tube, a dose and a stain. Gate for gate the constructs sat at comparable intensity: across the twenty arm × gate combinations the AQP9-to-GFP-Tubulin intensity ratio had a median of 1.41 (range 0.74 to 7.22) and the AQP3-to-GFP-Tubulin ratio a median of 1.48 (range 0.67 to 5.36). The gradient in Fig. 6 therefore compares cells carrying similar amounts of protein and differing in which protein they carry, and the amount carried is quantified rather than assumed.

## 4. Discussion

Most of an ingested inorganic arsenic dose leaves the body as MMA(V) and DMA(V) rather than as the parent compound^4^, and which transporters admit those metabolites, and what follows for the genome when they do, had not been established in a mammalian cell. We show that AQP9 and AQP10 confer susceptibility to DMA(V)-induced DNA damage in human cells, that AQP3 and AQP7 do not, that AQP9 alone remains active with MMA(V), and that the size of the effect is set by the host cell as much as by the channel. That damage scales with the amount of AQP9 a cell carries, and that GFP-Tubulin shows no such gradient, is the observation that ties the ranking to the channel rather than to transfection.

These results are consistent with the transport literature and reach an endpoint that literature does not. Human AQP9 in oocytes carries both MMA(V) and DMA(V)^20^, aquaglyceroporins carry MMA(III)^19^, and a bacterial aquaglyceroporin carries trivalent and pentavalent methylarsenicals alike^21^. Those studies measure flux; the 16.96-point difference reported here is what that flux costs the genome. It sits within an AQP9 literature in which the channel governs arsenic uptake and sensitivity in human cells^33,34^ and in immortalized human bladder epithelium^35^. One caveat runs the other way: aquaglyceroporins are bidirectional, and AQP9-null mice clear arsenic poorly and are more rather than less sensitive to it^36^, so a channel that raises damage in a cultured cell may mediate excretion in an intact animal.

AQP10 has almost no precedent. It has not been tested with a methylated arsenical in any system, and no genotoxic endpoint has been reported for it; the two prior measurements are an oocyte comparison in which it conducted little or no As(OH)_3_^22^ and an RNAi experiment in Caco-2 cells that changed As(III) uptake^23^. The gap has a structural cause: mouse Aqp10 is a pseudogene^25^, so the channel cannot appear in a rodent model however carefully that model is run. Human AQP10 is a functional glycerol channel^26^, so there was never a reason to expect it inert. That it conducted little or no inorganic arsenite yet gave the second largest DMA(V) effect measured here is the clearest single indication that permeability to an inorganic arsenical does not predict permeability to its methylated metabolites.

AQP3 makes the same point from the other side. The same construct in the same vector confers arsenite susceptibility in the companion study^27^, and here it left both methylated species unchanged. A channel that admits inorganic arsenite therefore does not detectably deliver the pentavalent methylated metabolites of that same arsenical to the genome. AQP7 sharpens it further: it conducts arsenite in oocytes^15^ and was inert to slightly negative for both methylated species here. Arsenical permeability is a narrow and separable property of an individual aquaglyceroporin, and it can differ between an inorganic species and its own metabolites within one channel.

Read across the four species we have now tested in this assay, genotoxic potency and aquaporin dependence do not follow the same order. In untransfected cells, arsenite raised the γH2AX-high fraction at 1 µM ^27^, As(V) needed 20 µM to move MRC5-SV40 cells and did not move HEK293T cells at all; and the pentavalent methylated species acted only in the millimolar range. That ordering follows membrane permeability, since uptake of trivalent arsenicals exceeds that of their pentavalent counterparts by orders of magnitude and correlates with toxicity^14^, and the species use different routes: arsenite through aquaglyceroporins^15^, As(V) through phosphate transporters^37^, and MMA(V) and DMA(V) through AQP9 at rates that rise as pH falls^20^. Our results divide along the same line, with the aquaporin-dependent increment confined to DMA(V) and, weakly, to MMA(V) through AQP9. In vivo a cell meets the whole series rather than one species, since ingested iAs is reduced to arsenite and methylated through MMA(V), MMA(III) and DMA(V), and DMA(V) is the arsenical most abundant in urine^4^. Aquaglyceroporin expression lowers the dose at which the least potent member of that series becomes genotoxic.

The present data extend rather than overturn the earlier comet screen of eight arsenicals in TK6 cells, in which MMA(V) and DMA(V) were inactive^9^. Three features of that design account for the divergent calls, and each point toward the same explanation. First, the endpoints differ: the comet assay detects strand breaks present at lysis, whereas γH2AX reports the damage response that follows. Second, the cell types differ in methylation and efflux capacity, which for arsenicals sets internal dose^14^. Third, and most consequentially here, the two measurements resolve a population very differently. A comet experiment is conventionally scored on the order of 10^2^ nuclei per sample and reported as a mean tail measure, while the flow cytometric assay used here reads 10^4^ to 10^6^ events per well and returns the whole γH2AX distribution rather than its average. A response carried by a minority of cells must therefore be large to move a comet mean but is read directly here as a shifted subpopulation. The present response takes exactly that form: the aquaporin differences widen rather than narrow as the γH2AX gate is relaxed (Figs. 2 to 5), and the excess damage is concentrated in the cells carrying most channel (Fig. 6). On this reading, the earlier null is what a population-averaged assay would be expected to return for a subpopulation effect of this magnitude, and the two datasets together delimit the conditions under which MMA(V) and DMA(V) activity becomes detectable. The concentration ranges also differ between the studies, a difference that cuts both ways.

That dose range is the principal limitation. The active range here is 0.5 to 10 mM, roughly a thousand-fold above the arsenite concentrations that produce comparable damage in the same assay and far above any concentration of a methylated metabolite measured in human tissue, so no claim about risk at ambient exposure follows from these data. What the experiment establishes is a mechanism and a ranking: at concentrations where these metabolites damage DNA, the damage is transporter-dependent, and which transporter a cell expresses changes it several-fold. Three further limitations bear on interpretation. No arsenical was measured inside a cell, so the uptake interpretation rests on published transport phenotypes rather than on a measurement made here; construct expression was not quantified against endogenous levels and no loss-of-function arm was run; and the trivalent methylated intermediates, which carry most of the genotoxic potency, were not tested.

Further work will measure intracellular arsenic speciation by ICP-MS across the four constructs, so that the ranking can be attached to an internal dose rather than inferred from one; will extend the panel to MMA(III) and DMA(III); and will test whether endogenous AQP9 and AQP10 levels predict methylated-arsenical sensitivity in primary human urothelium, the tissue in which DMA(V) is a demonstrated carcinogen.

## 5. Conclusion

Aquaglyceroporin expression sets how much DNA damage the dominant human arsenic metabolites produce in a human cell. AQP9 and AQP10 confer susceptibility to DMA(V), AQP3 and AQP7 do not, AQP9 alone remains active with MMA(V), and the damage a cell takes rises with the amount of channel that cell carries. The ranking does not follow permeability to inorganic arsenite, so a channel that admits an inorganic arsenical does not thereby admit its methylated metabolites, and the host cell scales the effect as strongly as the channel identity sets it. Because the active range is millimolar, these data establish a mechanism and a ranking rather than a statement about risk at ambient exposure. What they add to the assessment of methylated arsenicals is that genotoxic potency is a property of the arsenical and the transporter complement together rather than of the arsenical alone, and that a channel can act synergistically with a species weak enough to have been treated as a detoxification product.

## CRediT authorship contribution statement

**Shiwei Yin:** Investigation, Methodology, Validation, Writing – review & editing. **Maryam Vaziripour:** Investigation, Methodology, Writing – review & editing. **Bingru Feng:** Investigation, Resources, Writing – review & editing. **Jun Xia:** Conceptualization, Methodology, Software, Formal analysis, Data curation, Visualization, Supervision, Funding acquisition, Writing – original draft, Writing – review & editing.

## Data availability

All data supporting the conclusions of this study are presented in the article and its supplementary material. Further data and analysis code are available from the corresponding author upon reasonable request.

## Acknowledgements

We thank the Creighton University flow cytometry core facility for flow cytometry support.

---

**Supplementary Figure 1.**
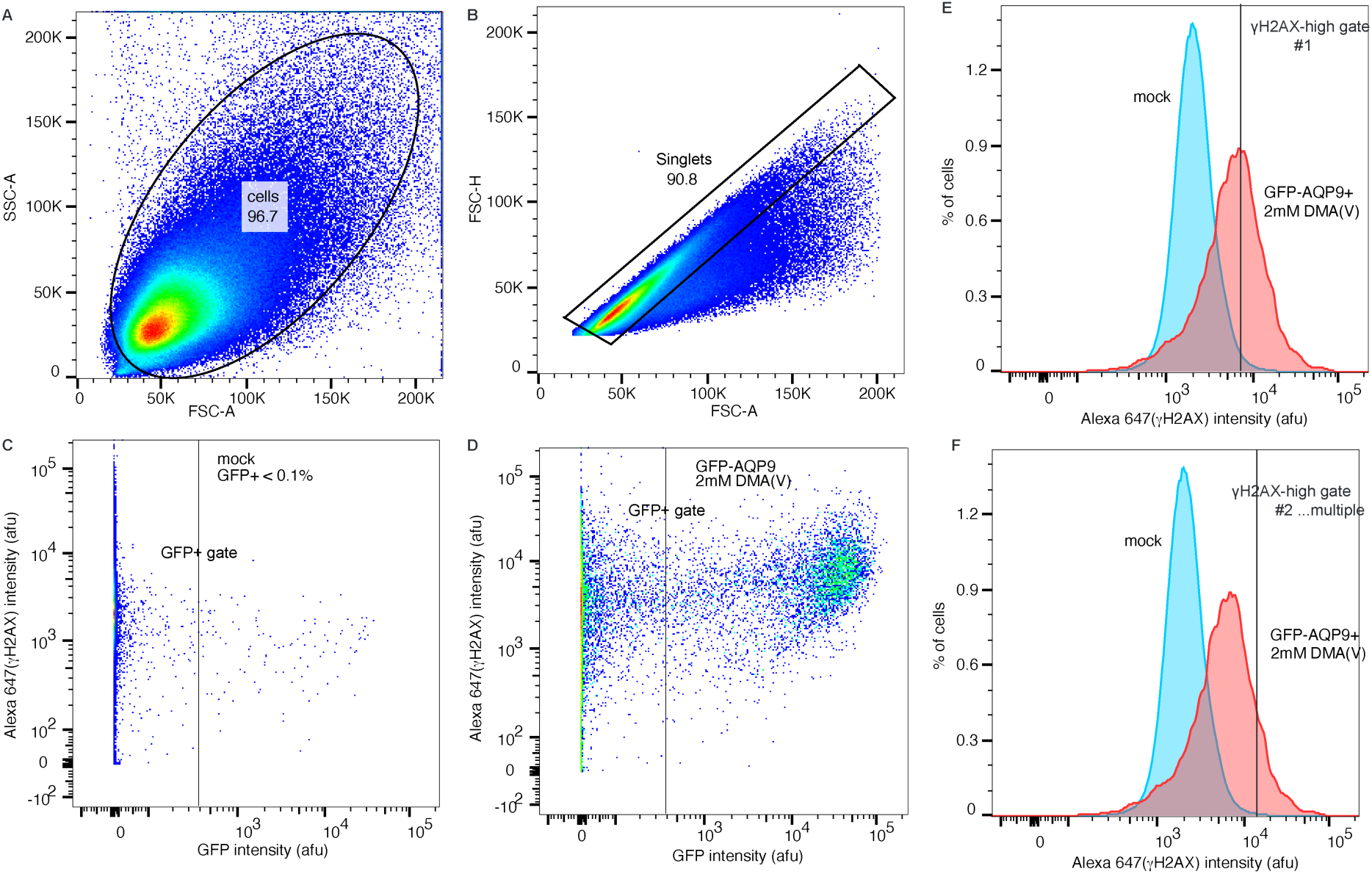
Gating strategy. Applied identically to all fcs files, shown on HEK293T, mock and AQP9 transfection with DMA(V) exposure. (**A**) Cells gate. (**B**) Singlets gate. (**C-D**) GFP-positive gate, the 99.9th percentile of the replicate’s own untransfected mock. (**E-F**) γH2AX-high gate, various, based on the replicate’s own untreated mock, applied to GFP-positive singlets.

**Supplementary Figure 2.**
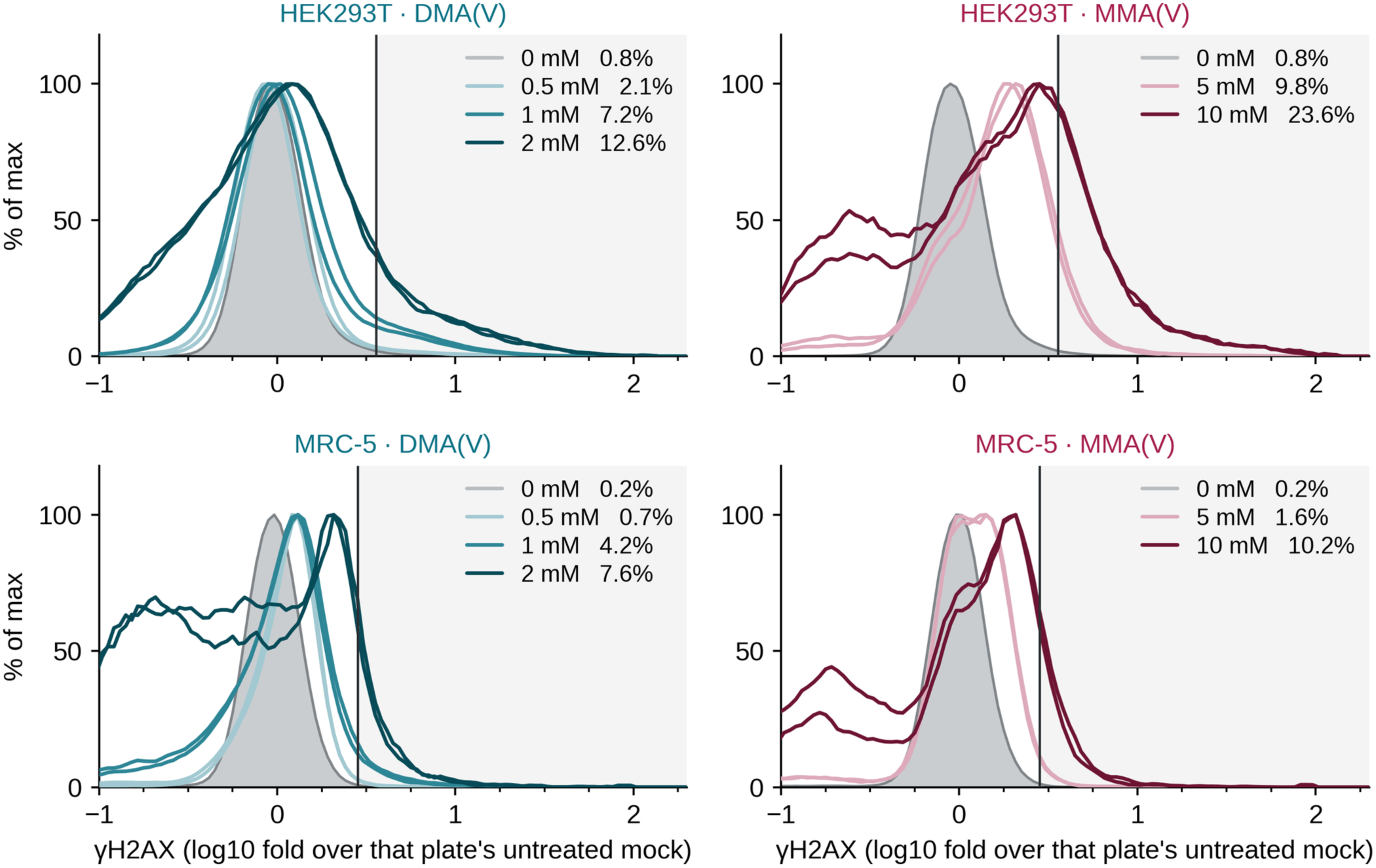
γH2AX distributions across the dose-finding ladders, overlaid. The data of Fig. 1C and 1D drawn as conventional overlays rather than as offset ridges, for readers who prefer to compare peak positions directly.

**Supplementary Figure 3.**
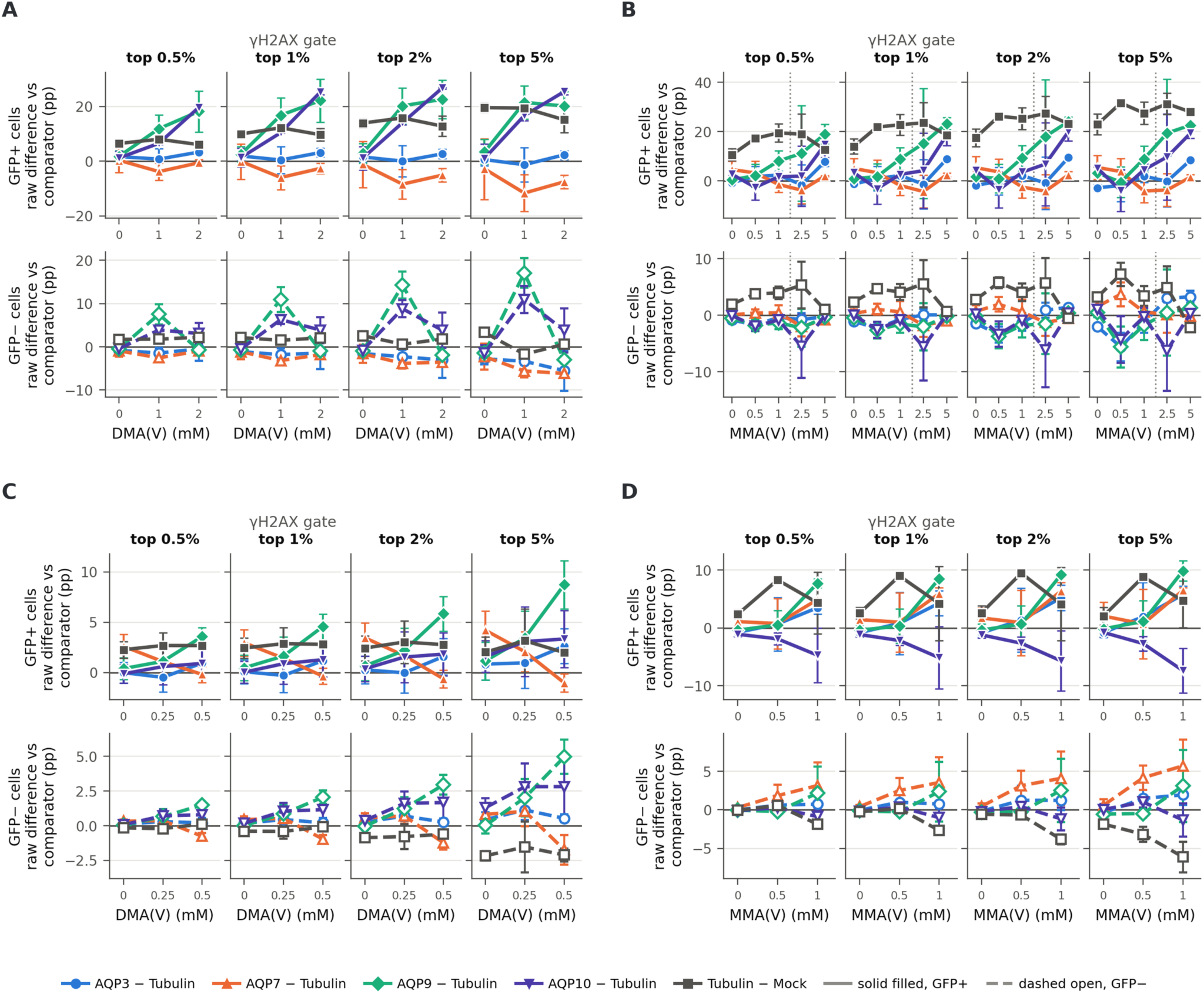
Raw paired differences at each dose. The contrasts of Figs. 2 to 5 resolved at each dose rather than pooled and given as the raw difference rather than the vehicle-subtracted one: the γH2AX-high percentage of an aquaporin well minus that of the GFP-Tubulin well beside it in the same replicate, and of GFP-Tubulin minus untransfected mock, without subtracting either construct’s own 0 mM well. The 0 mM column is therefore the baseline difference between two transfections before any treatment — the quantity vehicle subtraction removes — and the treated columns carry baseline plus treatment. (**A**) HEK293T DMA(V), (**B**) HEK293T MMA(V), (**C**) MRC5-SV40 DMA(V), (**D**) MRC5-SV40 MMA(V). Within each panel, columns are the four γH2AX gates, the upper row is GFP-positive cells and the lower row the GFP-negative cells of the same wells; mock is read on its GFP-negative population throughout. Mean ± SEM across replicates. The dotted rule in B marks where the dose axis is stitched from two replicate sets.

